# Hypoxia drives asymmetric envelope remodelling to promote membrane adaptation and antibiotic tolerance in mycobacteria

**DOI:** 10.64898/2026.09.12.751103

**Authors:** Lydia Mathew, Tzong-Hsien Lee, Marie-Isabel Aguilar, Shobhna Kapoor

## Abstract

*Mycobacterium tuberculosis* persists within the host by adapting to stressful microenvironments, with hypoxia representing a key clinically relevant challenge. However, how hypoxia alters mycobacterial membrane organization and function remains poorly understood. Using *Mycobacterium smegmatis* as a model, we combined membrane biophysics, layer-resolved lipidomics, and membrane-drug interaction analyses to characterize envelope adaptation under oxygen limitation. Hypoxia induced asymmetric remodelling of the cell envelope, with the inner membrane becoming more ordered and less hydrated, while the outer membrane exhibited increased fluidity and hydration. Lipidomic analyses revealed corresponding membrane-specific changes, including enrichment of saturated, tightly packed lipids in the inner membrane and unsaturated, inverted-conical lipids in the outer membrane. Functionally, hypoxic cells showed reduced envelope permeability and decreased susceptibility to rifampicin, while membrane partitioning studies suggested reduced rifampicin association with hypoxia-adapted inner membranes. We identify asymmetric envelope remodeling as a previously unrecognized principle of bacterial adaptation to environmental stress. Rather than responding as a uniform structure, the mycobacterial envelope undergoes coordinated but divergent remodeling in its constituent membrane layers during hypoxia. This spatially resolved response couples inner membrane rigidification with outer membrane fluidization, generating a functionally specialized envelope architecture that simultaneously preserves membrane integrity, modulates permeability, and reduces antibiotic association. These findings suggest that membrane asymmetry is not merely a structural feature of complex bacterial envelopes but an actively regulated adaptive strategy that links environmental sensing to stress resilience and antimicrobial tolerance

**Significance:** Most studies view bacterial membranes as responding uniformly to environmental stress. By resolving the inner and outer membranes separately, we show that hypoxia drives opposing remodelling programs within the mycobacterial envelope. This asymmetric adaptation alters membrane organization, permeability, and rifampicin interactions, revealing a previously unrecognized mechanism of physiological adaptation and antibiotic tolerance. The findings suggest that membrane asymmetry itself constitutes an important regulatory feature of pathogen survival under stress and may provide new opportunities for therapeutic intervention against persistent mycobacterial infections.

## Introduction

Tuberculosis (TB) persists as a major global health burden largely because of the remarkable adaptive capacity of the *Mycobacterium tuberculosis* (*Mtb*) bacterium within the host tissues. Rather than being rapidly eliminated, the bacillus establishes long-term survival inside granulomatous lesions—complex immune structures characterised by restricted vascularization, caseation, and limited oxygen ^1,2^. As granulomas mature, oxygen tension progressively declines, exposing bacilli to sustained hypoxic stress. This oxygen-limited environment promotes a non-replicative and likely drug-tolerant state that underlies prolonged chemotherapy and relapse ^3^. Hypoxia therefore represents a critical physiological environment during TB infection with an unknown spectrum of adaptive bacterial modifications.

Granulomas are both hypoxic and lipid-rich, containing foamy macrophages that accumulate host fatty acids and cholesterol, creating an environment of oxygen limitation and lipid abundance^4^. In response, *Mtb* activates the DosR dormancy regulon and regulators such as Rv0081, while remodelling its metabolism toward fatty acid β-oxidation and the glyoxylate shunt to maintain survival under restricted respiration^5,6^. Although these molecular adaptations are well characterized, hypoxia has been examined predominantly through transcriptional and metabolic frameworks. Yet oxygen limitation inherently challenges processes embedded within the membrane. The electron transport chain resides in the inner membrane, where proton motive force generation and redox balancing occur. Reduced oxygen availability constrains respiratory flux, potentially destabilizing membrane potential, altering lipid precursor availability, and perturbing lipid–protein interactions^7^.

Thus, beyond metabolic rewiring, hypoxia imposes a fundamental biophysical stress on the mycobacterial cell membrane envelope. This is further aggravated due to the most structurally complex and unique lipids that typify mycobacterial species. The cell envelope of *M. tuberculosis* is a multilayered structure comprising an inner membrane (IM), a peptidoglycan–arabinogalactan core (PG-AG), and an outer membrane (OM). The inner membrane contains classical phospholipids such as phosphatidylethanolamine (PE), phosphatidylinositol (PI) and phosphatidyl-myo-inositol mannosides (PIMs) with varying degrees of headgroup mannosylation (PIM_1-6_). The outer membrane is enriched in noncovalently attached free lipids such as mycolic acids (MA) and their ester derivatives, including trehalose mono- and di-mycolate (TMM, TDM). This layer also contains other non-canonical lipids such as sulfolipids-1 (SL-1), phthiocerol dimycocerosate (PDIM) and di-acyl or poly-acyl trehalose (DAT/PAT). Collectively, the mycobacterial cell envelope comprises an array of vertically stratified unusual lipids with varied physicochemical properties^8^. This architecture, in turn, influences permeability, inherent drug resistance (drug accessibility through the cell envelope), redox homeostasis, and mechanical resilience^9^. Furthermore, as membrane composition dictates bilayer thickness, curvature, fluidity, and stiffness, alterations in lipids are expected to have direct mechanical consequences^10^. However, in what way hypoxia reshapes membrane organization and mechanics and eventually regulates lipid-mediated mycobacterial survival and drug tolerance across envelope compartments remains unknown.

In this work, we combine multiparametric lipidomic profiling, microbiology, and membrane biophysics to decipher the role of hypoxia in modulating bacterial membranes, the underlying molecular mechanism and the ensuing consequence on membrane-centric antibiotic tolerance. We reveal that the mycobacterial cell envelope undergoes spatially choreographed fluidization and rigidification in a hypoxia-adapted growth-arrested yet viable state, inducing envelope softening. Delving deeper furnished that while the outer membrane becomes homogeneously fluidized, hypoxia induced heterogeneous induction of rigid/ordered and disordered/fluid domains within the inner membrane. Lipidome profiling demonstrated asymmetric redistribution of lipid species within the specific envelope layers under hypoxia, classified on the basis of lipid chain lengths, saturation levels and geometry. This altered distribution provided a structural basis for the observed membrane re-organization in both the cell envelope layers. Bacteria under hypoxic stress displayed attenuated permeability and tolerance to Rifampicin. Membrane-layer specific partitioning assays revealed that hypoxia-adapted IML functions as a highly restrictive secondary barrier that limits overall membrane permeabilization and drug association.

Our results identify asymmetric envelope modulation as a previously unknown paradigm of bacterial membrane adaptation. Under hypoxic stress, the mycobacterial cell envelope does not behave as a single adaptive unit; instead, its constituent membrane layers undergo opposing biophysical and lipidomic trajectories that collectively optimize survival and persistence upon antibiotic exposure. This spatially resolved response couples inner membrane rigidification with outer membrane fluidization, generating a coordinated architecture that reduces permeability and attenuates rifampicin-membrane interactions. These findings expand current models of bacterial stress adaptation beyond bulk lipid remodelling and establish membrane asymmetry as an emergent determinant of drug tolerance. Given the prevalence of multilayered envelopes among clinically important actinomycetes and other bacterial pathogens, asymmetric membrane remodelling may represent a broadly conserved mechanism linking environmental sensing to antimicrobial resilience.

## Results and Discussion

### Hypoxia induces a growth-arrested state with a soft cell outer envelope

The seminal work of Wayne and Hayes demonstrated that gradual oxygen depletion in a sealed culture system drives mycobacteria into defined stages of nonreplicating persistence, a process that can be visually monitored by the progressive fading and eventual decolourisation of methylene blue as oxygen levels decline^3^.

*Mycobacterium smegmatis* (*Msm*) was employed as a model for *Mtb*, given the substantial overlap in the lipidomes of both species as well as conserved metabolic pathways^11,12^. We observed a visible reduction in dye intensity between 10–15 h, with complete decolourisation by 25–30 h, confirming progressive oxygen depletion within the system^3^ **(Fig. S1).** Concomitant with oxygen decline, hypoxic cultures exhibited marked growth attenuation, reaching only ∼0.4 OD_₆₀₀_ compared to ∼4.2–4.3 OD_₆₀₀_ under normoxic conditions (**Fig. 1A, Fig. S2**). These observations confirm a successful establishment of a hypoxia-induced, slow-growing physiological state, as previously reported, suitable for subsequent nanomechanical and membrane-level analyses. Resazurin-based viability measurements further corroborated these findings. Normoxic cells displayed a progressive increase in metabolic activity over time, consistent with active proliferation (**Fig. 1B**). Conversely, hypoxic cells maintained significantly lower metabolic activity throughout the experiment, with only a modest increase at later time points (**Fig. 1B**). Together, these data confirm that oxygen depletion induces substantial growth restriction and metabolic downregulation in *Msm*^3^. The hypoxic model was functionally validated using metronidazole, a drug expected to exhibit enhanced activity specifically under oxygen-limited conditions. Propidium iodide (PI), which is generally excluded by cells with intact membranes but penetrates membrane-compromised cells (caused by cytotoxicity due to drug treatment), was used as a readout of cellular membrane integrity. Following metronidazole treatment, *Msm* demonstrated a clear dose-dependent increase in the proportion of PI-positive cells, whereas normoxic bacteria exhibited comparatively minimal PI uptake across the same concentration range. At the highest concentration tested (6400 µg/mL), approximately 37 % of hypoxic cells were PI-positive. This enhanced response under hypoxia provides functional evidence that the oxygen-limited state was successfully established and that the bacterial population exhibited the expected increased susceptibility to metronidazole under hypoxic conditions (**Fig 1C**). Next, using atomic force microscopy-based (AFM) nanoindentation^13^, we measured the mechanical properties of the bacterial cell envelope (i.e., stiffness/elastic modulus) under this state (**Fig. 1D-E**). Quantitative elastic modulus measurements revealed a significant reduction in stiffness in hypoxic bacteria compared to a normoxic control. Normoxic cells exhibited a broad distribution of higher modulus values, whereas hypoxic cells showed a distinct shift toward lower modulus values. Statistical analysis demonstrated a highly significant decrease, indicating that hypoxia markedly softens the bacterial outer layer of the envelope^14,15^.

**Figure 1.**
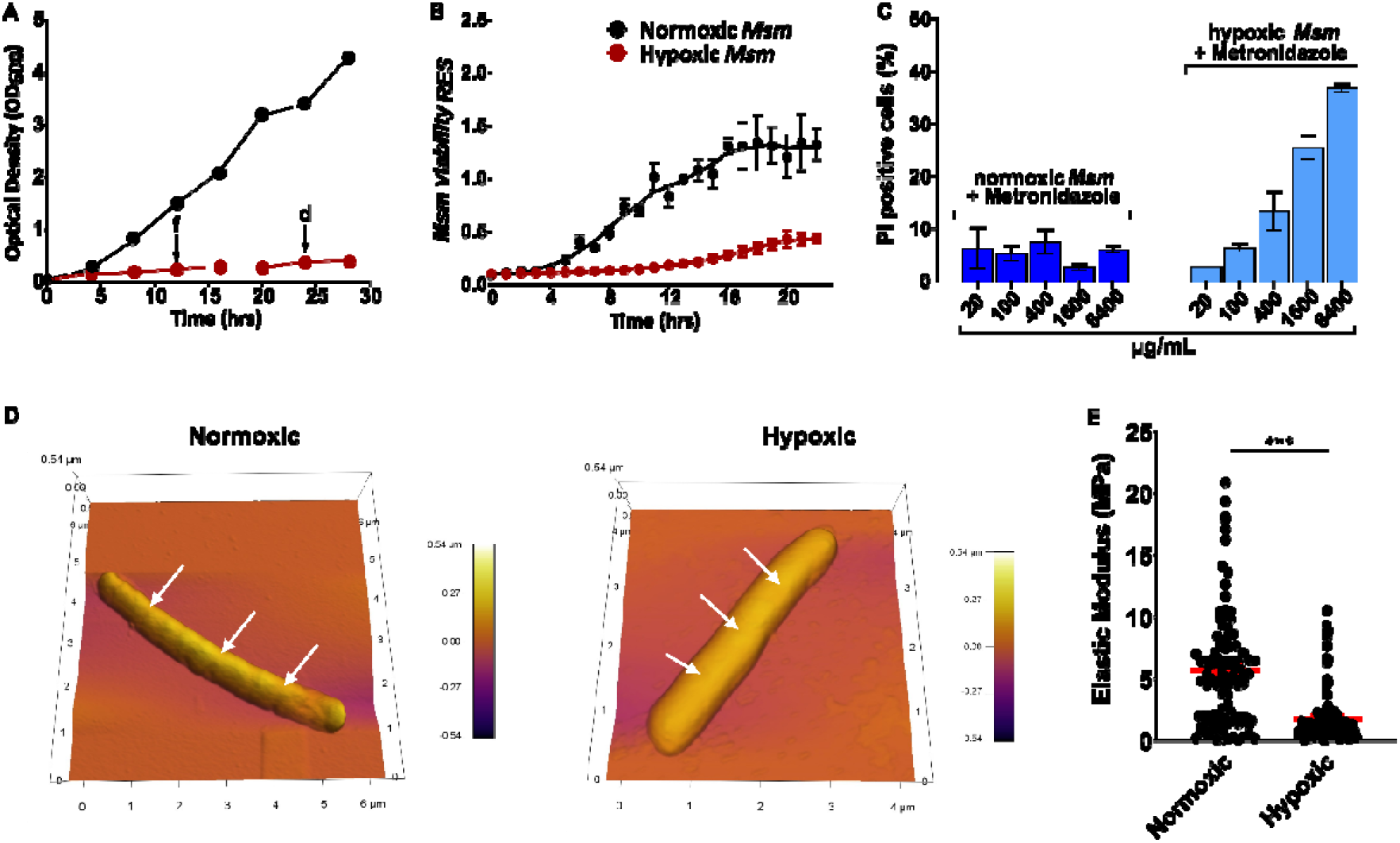
Establishment of hypoxic conditions and growth characterization of *Mycobacterium smegmatis*. (A) Growth kinetics of *Mycobacterium smegmatis* cultured under normoxic and hypoxic conditions measured by optical density at 600 nm (OD_₆₀₀_). Hypoxic cultures exhibited markedly attenuated growth compared to normoxic controls, reaching ∼0.4 OD_₆₀₀_ by 28–30 h, indicative of a slow-growing, dormancy-like state. Arrows indicate progressive fading (f) and complete decolourization (d) of methylene blue corresponding to oxygen depletion within the sealed culture system. (B) Resazurin-based viability assay showing metabolic activity of normoxic and hypoxic cultures over time. (C) Metronidazole treatment reveals enhanced, dose-dependent bacterial viability under hypoxia, functionally validating the hypoxic state. (D) Representative AFM topographic and three-dimensional reconstructions of M. smegmatis cultured under normoxic and hypoxic conditions, and white arrows indicate spots at which elastic modulus was measured. (E) Quantification of bacterial elastic modulus derived from AFM nanoindentation measurements. Red bars represent mean values. Data represent mean ± SEM. (Statistical significance was determined using appropriate statistical analysis (***p < 0.001)

### Asymmetric biophysical modulation of the inner and outer layers of the cell envelope under hypoxia

The mycobacterial cell wall consists of an outer and inner membrane layer^16^. Thus, to delineate the mechanistic basis of altered nanomechanics, lipids from both these layers (referred to hereafter as the outer and inner membrane lipids (OML or IML)) were extracted from bacteria cultured under both normoxic and hypoxic conditions using previously reported methods^16,17^. Large unilamellar vesicles (LUVs) were generated from these protein-free lipid extracts, and membrane order was assessed using Laurdan spectroscopy. Laurdan, being a solvatochromic probe, reports on the membrane microenvironment polarity by exhibiting a red-shifted emission in a more disordered/polar state^18^. The same is represented as a ratio metric parameter called generalized polarization (GP), with high GP indicative of a highly ordered membrane state and vis-a-vis. Biophysical characterization revealed a striking asymmetric response. Under hypoxia, IML exhibited increased rigidity/membrane ordering in a temperature-dependent manner with a lower sensitivity to temperature (between 20 °C and 40 °C; lower slope change) (**Fig. 2A**). In contrast, OML displayed increased membrane disorder in the hypoxic state compared to normoxia at all temperatures, with marginally higher sensitivity to temperature (**Fig. 2B**). This opposing behaviour indicates compartment-specific membrane adaptation rather than uniform softening of the entire envelope. Importantly, the magnitude of change under hypoxia was much larger in OML than in IML, suggesting that the increased outer membrane disorder most likely contributes to the reduced whole-cell elastic modulus (**Fig. 1D**). Simultaneously, inner membrane rigidification may serve to preserve essential membrane-associated processes during metabolic slowdown (**Fig. 1B**) ^14,19^.

**Figure 2:**
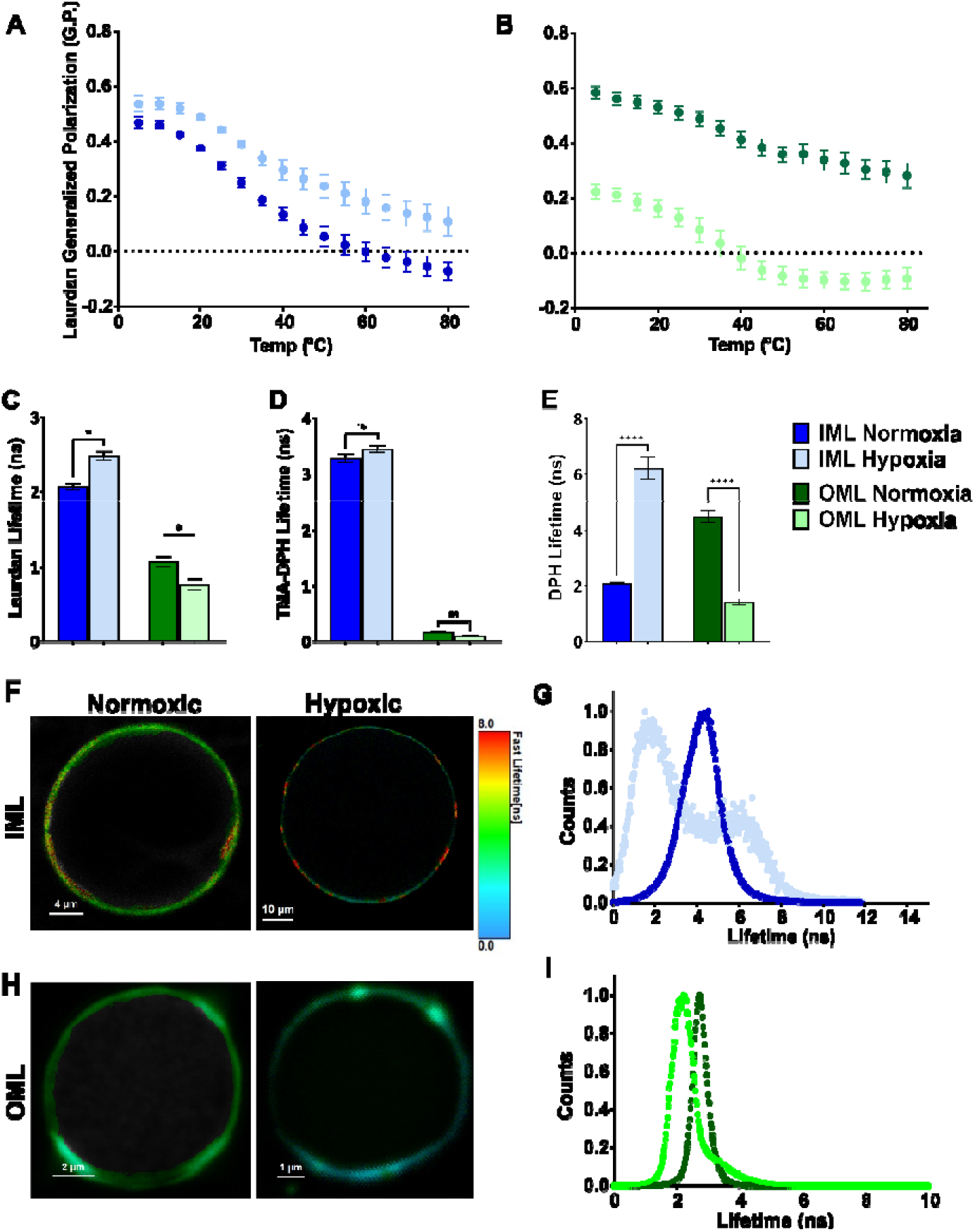
Asymmetric membrane remodelling under hypoxia revealed by Laurdan and fluorescence lifetime analyses. (A–B) Laurdan generalized polarization (G.P.) measurements of liposomes derived from inner membrane lipids (IML) and outer membrane lipids (OML). (C–E) TCSPC fluorescence lifetime measurements using Laurdan, TMA-DPH, and DPH probes. These findings demonstrate depth-dependent and asymmetric remodelling of the inner and outer membranes under oxygen limitation. (F–I) Representative Laurdan FLIM images and lifetime distributions of IML- and OML-derived GUVs. Data are mean ± SEM; *p < 0.05, ****p < 0.0001, ns, not significant.

To complement GP analysis, fluorescence lifetime measurements in bulk were performed using time-correlated single-photon counting (TCSPC) with Laurdan, TMA-DPH, and DPH probes targeting distinct membrane depths (**Fig. 2C-E**). These measurements allowed quantitative assessment of lipid packing at the headgroup interface (Laurdan), near the glycerol backbone region (TMA-DPH), or within the hydrophobic acyl chain core (DPH)^20,21^. In IML, hypoxia induced a significant increase in fluorescence lifetime compared to normoxia, indicating reduced interfacial hydration and enhanced lipid packing, supporting the GP analysis. Likewise, OM-derived membranes exhibited a significant decrease in Laurdan lifetime under hypoxia, reflecting increased headgroup mobility and greater water penetration. DPH lifetime demonstrated pronounced and statistically significant changes in the acyl chain region. Hypoxic IML exhibited a marked increase in DPH lifetime relative to the normoxic state, consistent with tighter acyl chain packing and reduced dynamic quenching within the hydrophobic core as well. In contrast, hypoxic OML displayed a substantial decrease in DPH lifetime compared to normoxic OML, indicating enhanced tail disorder and increased membrane fluidity. These data indicate that the effect of hypoxia on membrane dynamics spans from the head region to the hydrophobic core. Notably, TMA-DPH lifetimes showed minimal changes between normoxic and hypoxic conditions in both IML and OML fractions. The absence of significant differences suggests that structural alterations at the intermediate glycerol backbone region are modest compared to changes observed at the headgroup and deep hydrophobic core^22^. Alternatively, TMA-DPH may exhibit limited sensitivity to subtle interfacial rearrangements^21^. Collectively, it can be stated that hypoxia-induced remodelling is not uniformly distributed across the bilayer thickness but is spatially selective, predominantly affecting the bilayer surface and deep acyl chain regions, while preserving the interfacial transition zone.

Next, to visualize the above alterations in the membrane’s lateral organization at a single vesicle level, Laurdan fluorescence lifetime imaging microscopy (FLIM) was performed ^23^. This enabled us to elucidate the spatially induced effects of hypoxia on IML and OML compared to the above ensemble measurements. Giant unilamellar vesicles (GUVs) were generated from isolated IML and OML, and representative FLIM images revealed distinct lifetime distributions between normoxic and hypoxic conditions for both membrane systems. Hypoxic stress induced lateral membrane re-organization in IML, accompanied by the emergence of more ordered membrane domains (red domains) compared to the normoxic state (**Fig. 2F**). Quantitative analysis demonstrated a bimodal distribution of lifetime population (**Fig. 2G**), indicating the presence of discernible ordered and disordered domains simultaneously instead of a homogeneous (unimodal) distribution as seen in the normoxic state. The ordered domain, characterized by a higher lifetime (∼6.12 ns), accounted for ∼ 38.2% of the total population, reflecting domains of increased lipid packing, i.e., localized enhanced rigidity within inner membrane regions under oxygen limitation. Whereas the disordered domains, with a shorter lifetime (∼1.54 ns), comprised ∼61.7% (**Fig. 2G).**

Usually, domain heterogeneity is characteristic of multi-component lipid membranes containing coexisting lipid environments with varying degrees of order and hydration^24^. The above data indicate that under hypoxia, this is further enhanced locally within IML, leading to a rewired membrane domain organization with at least two discernible lipid domains (**Fig. 2G**). This suggests that hypoxia likely induces local clustering of selective inner membrane lipids into distinct ordered and disordered domains. On the other hand, OML-derived GUVs displayed a significant decrease in Laurdan lifetime under hypoxia relative to normoxia (**Fig. 2H**). The leftward shift in lifetime distribution reflects reduced lipid packing (green-blue domains) and increased water penetration at the membrane interface, indicative of enhanced outer membrane fluidity (**Fig. 2I**). Unlike in IML, the fluidization in OML was rather uniform, suggestive of distinct laterally resolved remodelling in outer and inner membranes of the mycobacterial cell wall under hypoxia. Importantly, combined with AFM, all these findings suggest that the low elastic modulus in intact bacteria under hypoxia not only stems from the OML layer but could also have a contribution from the *de novo* generated disordered regions within the IML seen at the single vesicle level. Together, varied membrane biophysical investigations confirm asymmetric mycomembrane remodelling under hypoxia. This unprecedented compartment-specific and spatially-resolved lipid reorganization provides a unusual basis for the reduced whole-cell elastic modulus observed under hypoxic conditions. This underscores the strategic modulation of membrane architecture as part of the mycobacterial adaptive responses to intracellular environments encountered within the host.

### Layer-Resolved Lipidomics of Mycobacterial Envelope Under Hypoxia

To uncover the compositional and structural basis of the membrane remodelling, we performed a multi-parameteric lipidomics approach on the inner and outer membrane lipids. Following lipidomics, mycobacterial lipids were classified into six major classes: fatty acyls, glycerophospholipids, glycerolipids, saccharolipids, polyketides, and prenol to establish a global compositional framework before subclass-level analysis. Such lipid classification has been widely employed in mycobacteria^11,19,25^.

To determine whether these compositional changes generated distinct membrane-specific lipid states, principal component analysis (PCA) was performed separately on IML and OML lipidomes^26^. Both membrane fractions showed clear segregation between normoxic and hypoxic samples, indicating that hypoxia induces reproducible and well-defined changes. IML clustered distinctly along the principal component axis, suggesting coordinated alterations in lipid composition associated with membrane adaptation. Similarly, the OML exhibited strong separation between conditions, reflecting substantial remodelling of outer envelope-associated lipids. The distinct clustering patterns further support the notion that the IML and OML respond differently to hypoxic stress **(Fig. S3A-B)** ^6,19^.

Global lipidomic analysis, supported by heatmap visualization of lipid class abundance, identified glycerophospholipids as the most abundant lipid class in both the inner and outer membrane fractions under normoxic and hypoxic conditions, followed by fatty acyls and saccharolipids. Despite their predominance, glycerophospholipids underwent membrane-specific remodelling in response to hypoxia. Comparison between normoxic and hypoxic conditions demonstrated a significant increase in glycerophospholipids within the IML, accompanied by a concomitant decrease in fatty acyls, whereas the OML displayed the opposite trend, with a significant reduction in glycerophospholipids together with increased fatty acyls and saccharolipids (**Fig. 3A-B**)

**Figure 3:**
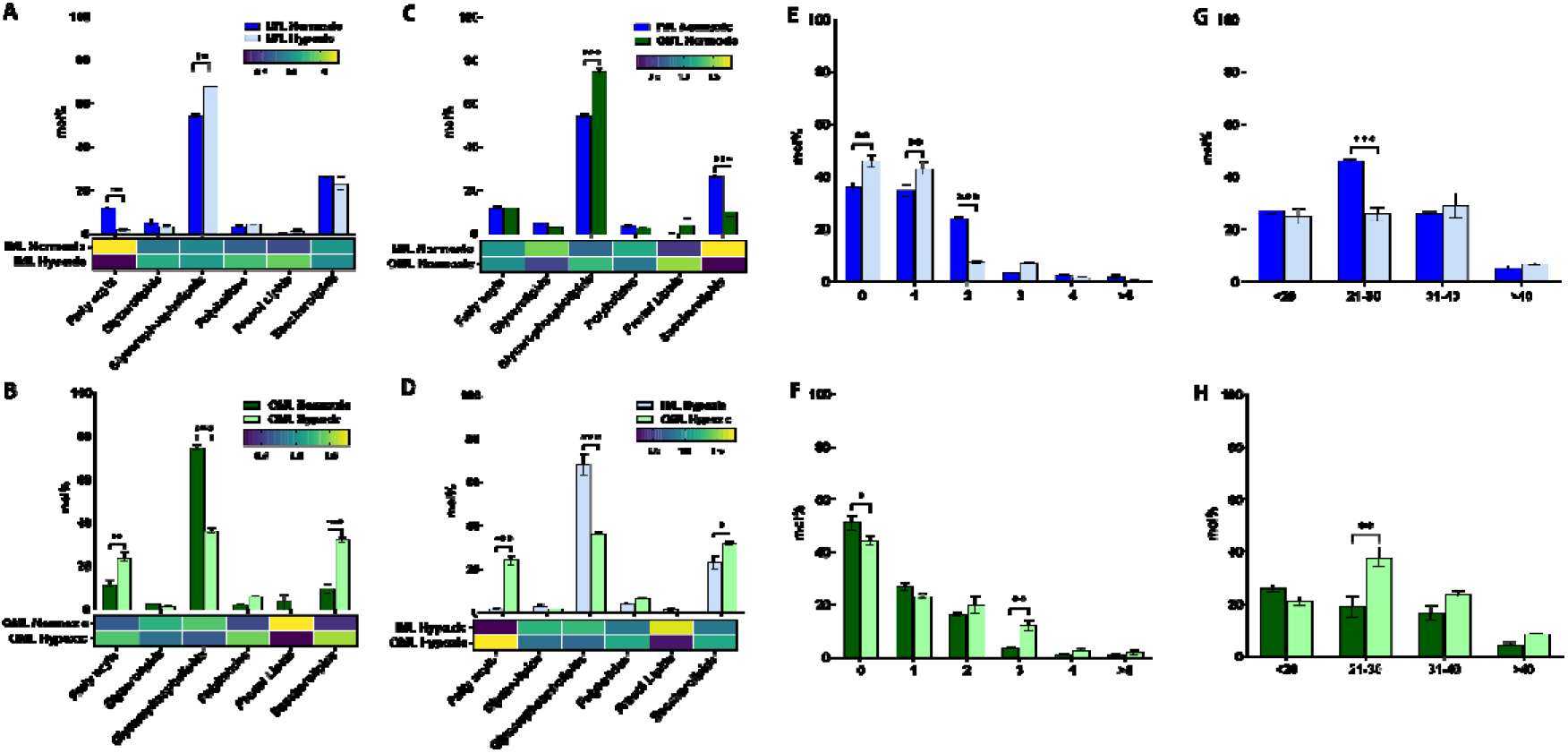
Lipidomic profiling of membrane lipids in the inner and outer membrane fractions of *Mycobacterium smegmatis* under normoxic and hypoxic conditions. Comparison of the relative abundance of major lipid classes between normoxic and hypoxic conditions in the (A) inner membrane lipid (IML) and (B) outer membrane lipid (OML) fractions, with the corresponding heatmaps showing relative abundance. Comparison of lipid class distribution between IML and OML under (C) normoxic and (D) hypoxic conditions, with corresponding heatmaps. (E, F) Overall distribution of lipid unsaturation levels in the IML and OML, respectively. (G, H) Overall distribution of lipid acyl chain lengths in the IML and OML, respectively. Data are presented as mean ± SEM.

Comparison of the two membrane layers within each physiological condition further revealed a pronounced lipid asymmetry. Under normoxic conditions, OML contained a higher proportion of glycerophospholipids than the IML, while the opposite was seen in hypoxic conditions (**Fig. 3C-D**). Notably, hypoxia further accentuated this asymmetry by increasing glycerophospholipid abundance in the IML while decreasing it in the OML. Collectively, these findings demonstrate that, although glycerophospholipids remain the dominant lipid class in both membrane fractions, hypoxia drives membrane layer-specific lipid remodelling, resulting in enhanced compositional asymmetry between the inner and outer membranes. Similarly, saccharolipids decreased in OML under normoxic conditions, while in hypoxic conditions, their levels increased more than in IML. Furthermore, fatty acyls accumulated in OML under hypoxia but not in normoxic conditions. These selective lipidomic alterations correlate with the contrasting biophysical properties observed under hypoxic conditions, wherein the IML exhibited increased membrane order and reduced hydration. At the same time, the OML displayed an enhanced and uniform membrane fluidization. (**Fig. 3C-D**).

Next, the overall distribution of lipid unsaturation and acyl chain lengths within each membrane fraction was analysed (**Fig. 3E-H**), as these parameters strongly influence membrane packing, fluidity and rigidity ^10^. Within the IML, hypoxic adaptation resulted in a significant increase in the abundance of saturated and mono-unsaturated lipid species, accompanied by a significant reduction in di-unsaturated lipids compared to the normoxic condition (**Fig. 3E**). This increase in saturated and mono-unsaturated lipids can contribute to both the ordered and *de novo* fluidic domains observed within IML hypoxic GUVs (**Fig. 2F**). In contrast, the OML exhibited significant decrease in saturated lipids and a corresponding increase in tri-unsaturated lipid species under hypoxia (**Fig. 3F**), underlying the observed uniform membrane fluidization (**Fig. 2H**). Comparison of the two membrane layers within each physiological condition also highlighted a pronounced asymmetry in unsaturation profiles. Under normoxic conditions, the OML was enriched in saturated lipid species, whereas the IML contained relatively higher proportions of mono- and di-unsaturated lipids. Hypoxia further accentuated this asymmetry, with the IML shifting towards a more saturated lipid profile, while the OML displayed a relative enrichment of tri-unsaturated species (**Fig S4A-B**).

Analysis of the overall acyl chain length distribution also revealed discrete membrane-specific remodelling under hypoxia. Within the IML, hypoxia resulted in a significant reduction in the abundance of the intermediate chain length lipids, i.e., 21–30 carbon atoms, whereas the proportions of <20, 31–40, and >40 carbon lipids remained largely unchanged (**Fig. 3G**). In contrast, the OML exhibited the opposite trend, with a significant enrichment of the same category of 21–30 carbon lipid species with no significant changes in the remaining acyl chain length populations (**Fig 3H)**. These coordinated changes provide a structural basis for the biophysical properties observed under hypoxic conditions. In IML, enrichment of saturated lipid species together with a reduction in intermediate-chain (21–30 carbon) lipids correlates with increased membrane order and reduced hydration. In contrast, the OML exhibits a reduction of saturated lipids and enrichment of tri-unsaturated and intermediate-chain (21–30 carbon) lipid species, consistent with enhanced membrane fluidity and increased membrane hydration. Comparison of the two membrane fractions within each condition revealed that hypoxia reduced the abundance of 21–30 carbon lipid species in the IML, the same lipid population increased significantly in the OML, demonstrating that the two membrane layers undergo opposing acyl chain modulation in response to oxygen limitation (**Fig. 3G-H**).

All these findings cement the proposition that hypoxic adaptation does not involve a uniform restructuring of membrane lipids but rather promotes a differential remodelling of the inner and outer membrane layers, thereby enhancing the intrinsic asymmetry of the mycobacterial cell envelope.

### Multi-parametric lipidome profiling reveals specific changes in molecular geometry across the layers

Expanding further on the lipidomic changes, the lipids were subsequently categorized according to their molecular geometry into conical, inverted conical, and cylindrical species (**Fig. 4-5**). As lipid shape directly governs membrane packing defects, hydration, and distribution, this analysis provided the mechanistic insight into the biophysical changes. Recent studies further emphasise that collective lipid shape distributions can profoundly regulate membrane mechanics, curvature adaptability, and stress-responsive membrane organization^25,27,28^. Global analysis demonstrated that inverted conical lipids constituted the predominant lipid geometry in both the inner membrane (IML) and outer membrane (OML), followed by conical and cylindrical lipids (**Fig. 4A**, **5A**). Conical lipids possess relatively small headgroups and bulky hydrophobic tails, generating negative intrinsic curvature stress and promoting localised packing defects within the bilayer^29,30^. Their accumulation is therefore expected to increase membrane flexibility and facilitate curvature adaptation under stress conditions. Cylindrical lipids favour planar bilayer organisation and tighter lateral packing due to the balanced cross-sectional areas of their headgroups and acyl tails^28,30^.

**Figure 4:**
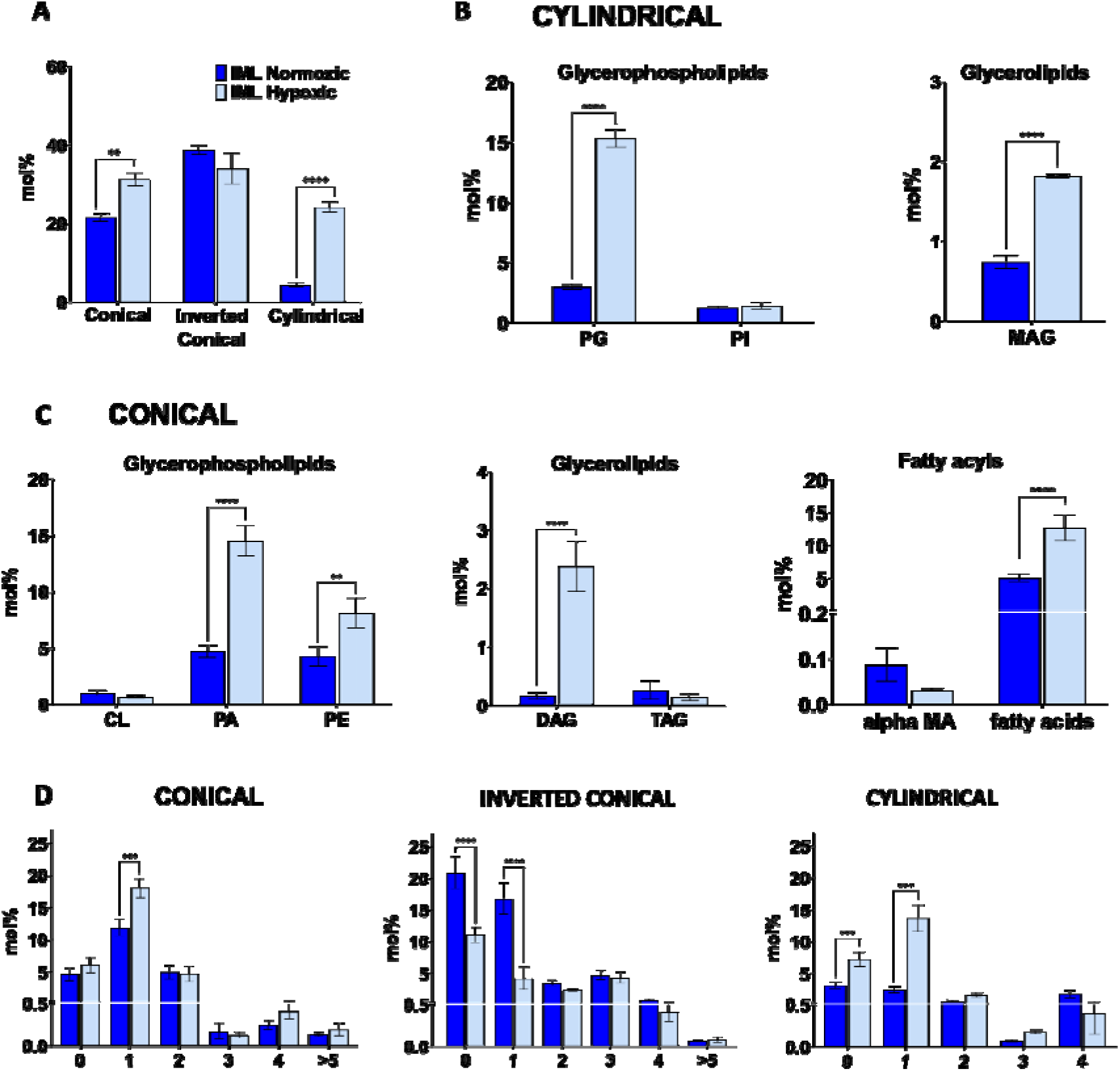
Hypoxia-induced remodelling of membrane curvature-associated lipid species in the inner membrane of *Mycobacterium smegmatis*. The relative abundance of lipid species categorized according to their intrinsic molecular geometry was determined from inner membrane (IM) lipidomic datasets under normoxic and hypoxic conditions. (A) Overall distribution of conical, inverted conical, and cylindrical lipid classes, among which conical and cylindrical lipid geometry categories showed changes. (B) & (C) Distribution of lipid species within cylindrical and conical geometric categories. (D) Distribution of lipid species based on levels of unsaturation or number of double bonds within conical and cylindrical lipid geometry categories. Lipid abundances are expressed as mol%, and bars represent mean ± SEM from 3 biological replicates. Statistical significance was determined using Student’s t-test (P < 0.05, **P** < 0.01, **P** < 0.001, **P** < 0.0001).

**Figure 5:**
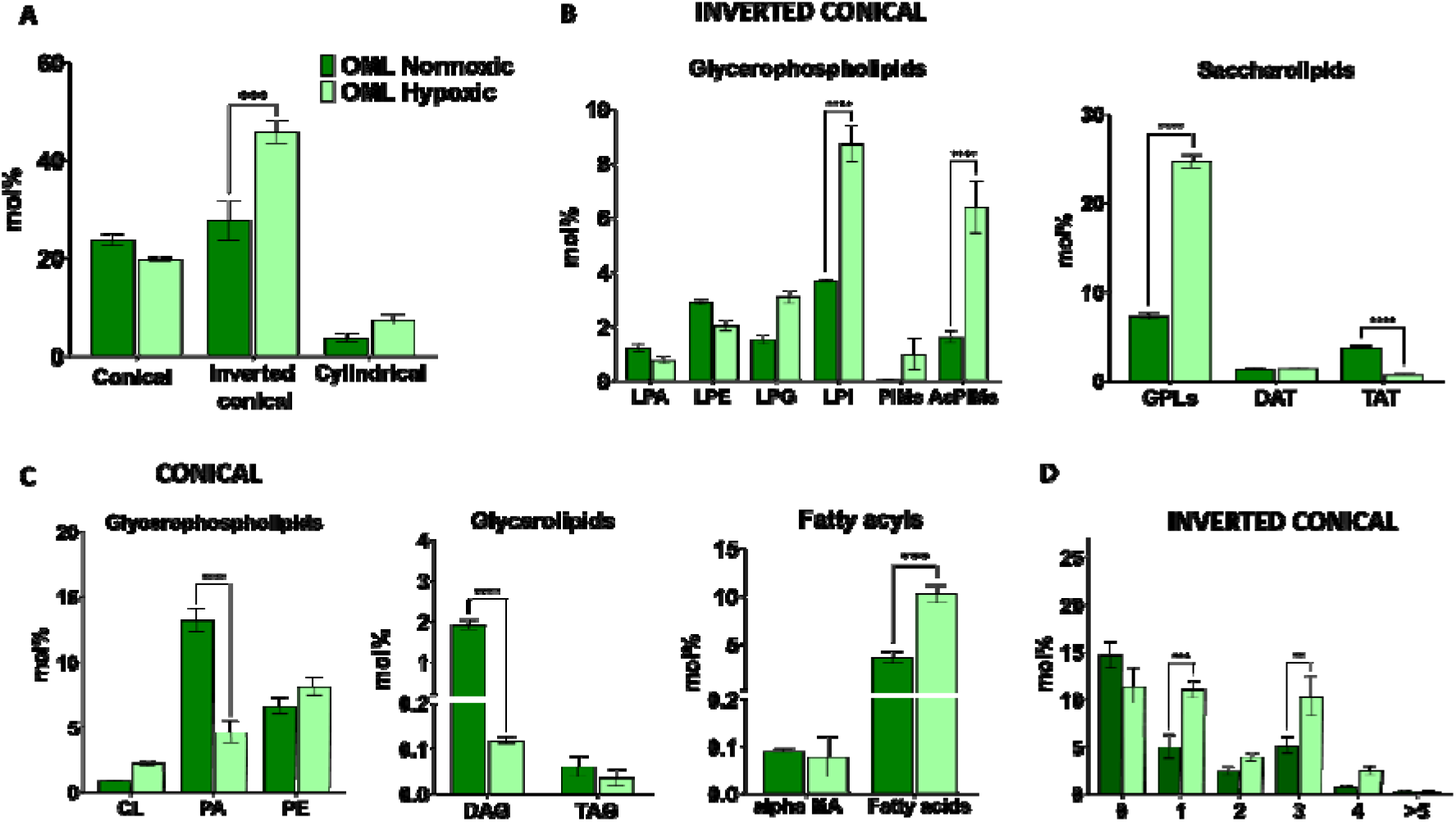
Hypoxia-induced remodelling of membrane curvature-associated lipid species in the outer membrane of *Mycobacterium smegmatis*. The relative abundance of lipid species categorized according to their intrinsic molecular geometry was analyzed from outer membrane (OM) lipidomic datasets obtained under normoxic and hypoxic conditions. (A) Overall proportions of conical, inverted conical, and cylindrical lipid classes, among which we saw a change within the inverted conical and conical lipid geometry categories. (B) & (C) Distribution of lipid species within the inverted conical and conical lipid geometric categories. (D) Distribution of lipid species based on levels of unsaturation or number of double bonds within the inverted conical lipid geometry category. Lipid abundances are represented as mol%, with bars indicating mean ± SEM from 3 biological replicates. Statistical significance was assessed using Student’s t-test (P < 0.05, **P** < 0.01, **P** < 0.001, **P** < 0.0001).

To determine the lipid species that are responsible for the altered membrane behaviour and geometry under hypoxia, individual lipid subclasses were examined within each geometry. Overall, hypoxia increased the conical and cylindrical lipid species abundance within IML, while the inverted conical lipids remained unchanged (**Fig. 4A**). Enrichment of conical lipids was primarily driven by increased phosphatidic acid (PA), diacylglycerol (DAG), and free fatty acyl species across different lipid classes. At the same time, cylindrical lipids such as phosphatidylglycerol (PG) and monoacylglycerols (MAG) belonging to glycerophospholipids and glycerolipids, respectively, were also enriched (**Fig. 4A-B**).

Since membrane packing is also influenced by the levels of unsaturation, their distribution within each geometric category was also examined. Hypoxia induced specific changes in the unsaturation profiles of conical, inverted conical and cylindrical lipids rather than a uniform alteration across these classes (**Fig. 4C**). Monounsaturated species increased significantly among the conical and cylindrical lipids, whereas saturated lipids within the inverted conical lipid category decreased. These could contribute to the fluidic or lower lifetime domains present in the IML hypoxic GUVs (**Fig. 2F**). Concurrently, a decrease in the abundance of monounsaturated inverted conical lipids, together with an increase in cylindrical saturated lipids, likely contributes to the formation of more ordered and rigid domains. This observation is also consistent with the increased membrane order observed in bulk fluorescence measurements including Laurdan GP (Fig. 2A) and lifetime (Fig. 2C) and DPH lifetime (Fig. 2E) analysis, and is further supported by the rigid regions (yellow-red domains) detected in hypoxic IML GUVs (Fig. 2G).

In OML, hypoxia also induced a marked remodelling of lipid geometry, characterised by reduced levels of cylindrical lipids overall, and a significant increase in the inverted conical lipids (**Fig. 5A**). These changes indicate a shift towards lipid species possessing relatively larger hydrophobic tails, consistent with membrane flexibility seen under hypoxic stress.

Inverted conical lipids were enriched with lysophosphatidylinositol (LPI), acylated phosphatidylinositol mannosides (AcPIMs), and glycopeptidolipids (GPLs). Inverted conical lipids contain relatively large headgroups compared to their acyl tails and preferentially promote positive curvature and increased interfacial spacing between lipid headgroups ^27,29^. Their accumulation is therefore expected to increase membrane hydration and reduce tight packing at the membrane surface. This interpretation is consistent with the altered hydration and rigidity profiles observed in fluorescence-based measurements. The overall abundance of conical lipids remained unchanged due to a concomitant increase of free fatty acid lipids balanced by the reduction of phosphatidic acid (PA) and diacylglycerol (DAG), thereby likely only locally (i.e., over small length scales) contributing to observed membrane properties (**Fig. 5B**). This also suggests that, though local lipid rewiring may happen under hypoxia, it is the collective behaviour of the lipids that undermines the observed biophysical changes in the respective mycobacterial membranes and could be a conserved response in mycobacterial species. Further analysis of unsaturation within inverted conical lipids showed a significant increase in both mono- and polyunsaturated lipid species under hypoxia (**Fig. 5C**). The greater number of double bonds introduces bends within the acyl chains, reducing lipid packing efficiency and weakening van der Waals interactions between neighbouring lipids. These highly unsaturated inverted conical lipids are expected to promote the formation of more fluid, disordered membrane regions in the outer membrane layer under hypoxia (**Fig. 5C**). These membrane-specific geometric adaptations provide a structural basis for the differential fluidity, rigidity, and hydration behaviours observed under hypoxic stress. Recent lipidomic–biophysical studies further demonstrate that coordinated redistribution of lipid geometries and saturation states can generate distinct membrane material properties under stress adaptation ^25^.

Collectively, these findings demonstrate that hypoxia induces coordinated asymmetric lipid remodelling across the mycobacterial envelope with distinct changes occurring in the outer and inner membrane layers. In the OML, enriched levels of unsaturated conical and inverted conical lipids are expected to reduce lateral lipid packing and enhance acyl chain flexibility. These compositional changes are consistent with increased membrane hydration and packing observed by Laurdan GP (Fig. 2B), lifetime (Fig. 2C), DPH lifetime (Fig. 2E) and FLIM imaging (Fig. 2H).

### Hypoxia remodels bacterial envelope permeability to modulate antibiotic tolerance and drug partitioning

Following biophysical and lipidomic characterization of the hypoxic-induced membrane perturbation, we next investigated whether these changes are associated with any altered envelope permeability. As DAPI can enter compromised membranes, its uptake was used readout to compare dye internalization within normoxic and hypoxic *Msm* states. Normoxic bacteria showed substantially higher DAPI fluorescence (∼67%) than hypoxic bacteria (∼32%), suggesting that reduced DAPI fluorescence in hypoxic cells reflects lower permeability of the envelope associated with this state. Next, the effect of hypoxia on permeability in the presence of known membrane permeabilizers, Triton X-100 and Lysozyme was investigated. Normoxic bacteria displayed increased DAPI uptake (∼88%) compared with the untreated control (∼67%), confirming that artificially induced membrane permeabilization enhances DAPI uptake. In contrast, reduced DAPI uptake was seen in hypoxic bacteria (∼55%) even in presence of membrane permeabilizers. Together, these results indicate two distinct effects: first, hypoxic bacteria exhibited a lower DAPI accessibility in the absence of any treatment, and secondly, the increase in DAPI uptake induced by known membrane permeabilizers was further attenuated under hypoxia.

Seeing that hypoxia alters the bacterial envelope permeability, we next examined whether this could have a functional consequence for the activity of commonly used antibiotic, Rifampicin. Normoxic *Msm* exhibited a pronounced dose-dependent increase in PI-positive cells, particularly at concentrations approaching and exceeding the MIC. In contrast, hypoxic *Msm* consistently displayed markedly attenuated % of PI positive cells, even at concentrations several-fold above the MIC. These findings suggest that hypoxia-induced alteration of envelope organization and permeability likely enhances the tolerance towards the antibiotic by likely reducing its entry and subsequent activity.

Next, we have previously shown that rifabutin (an analogue of rifampicin) partitions distinctly within mycobacterial inner and outer membranes and that the membrane properties are directly associated with its localization^31^. Thus, to delineate which membrane layer governed the attenuated antibiotic activity within intact bacteria (**Fig. 6B**), *in vitro* drug partitioning using extracted OML and IML liposomes obtained from normoxic and hypoxic *Msm* was performed. Drug partitioning was quantified spectrophotometrically by monitoring rifampicin distribution between aqueous and membrane-associated phases, and partition coefficients (Kp) together with log D values were calculated as indicators of membrane affinity^32,33^.

**Figure 6:**
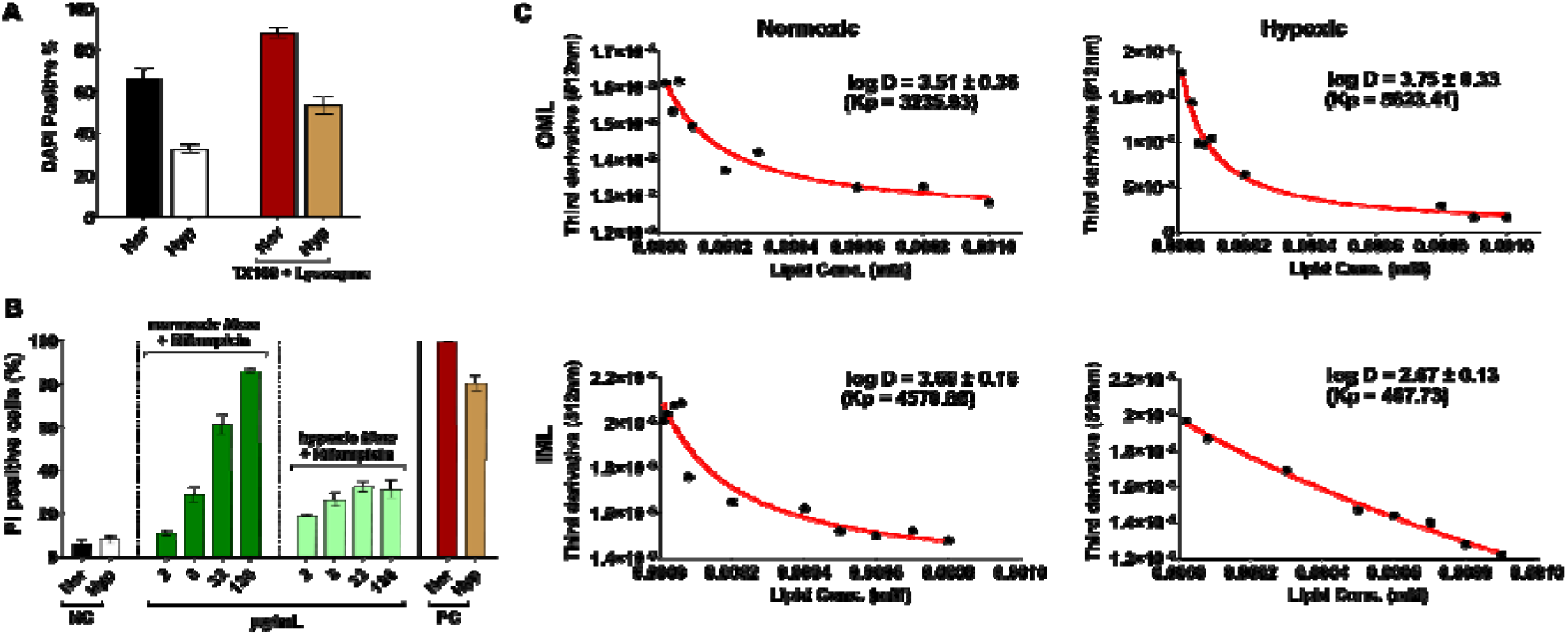
(A) DAPI uptake as a measure of bacterial envelope permeability in normoxic and hypoxic *Msm.* DAPI uptake was quantified in untreated *Msm* under normoxic and hypoxic conditions to assess basal membrane permeability. TX-100 and lysozyme were used as positive controls to induce bacterial envelope permeabilization and define the maximum DAPI uptake response under each condition. DAPI-positive cells were quantified by flow cytometry and expressed as the percentage of total bacterial cells. **(B) Propidium iodide (PI)-based flow cytometric analysis of viability in normoxic and hypoxic *Msm* following rifampicin treatment.** Hypoxic bacteria exhibited reduced PI uptake following rifampicin exposure compared to normoxic cells. Heat-killed bacteria were used as positive controls (PC). **(C) Rifampicin (10 µM) partitioning profiles in isolated IML and OML fractions from normoxic and hypoxic *Msm*.** Data represent mean ± SEM of 3 biological replicates. Data were fitted using a non-linear membrane partitioning model to determine log D values.

Despite the increased fluidic heterogeneity observed in OML under hypoxia, the overall membrane affinity of rifampicin remained relatively unchanged, indicating that OML remodelling alone is not sufficient to modulate drug interaction. In contrast, the IML under hypoxia showed a marked reduction in rifampicin partitioning compared with the normoxic condition (**Fig. 6C**), suggesting a strong correlation with the increased rigidity and ordered domain formation induced within IML under hypoxic stress. Lipidomic analyses further revealed enrichment of cylindrical and tightly packed lipid species together with reduced interfacial disorder in this membrane fraction. The resulting condensed membrane organisation likely limits rifampicin insertion and diffusion within the inner membrane bilayer thereby attenuating the overall drug activity within intact bacteria^32,33^. Collectively, hypoxia-adapted IML appears to function as a highly restrictive barrier that limits overall membrane permeabilization. The biophysical assays done on reconstituted liposomes from both the layers under normoxic and hypoxic states demonstrate that the remodelled lipid composition is sufficient to alter membrane properties and rifampicin partitioning.

Our study advances a conceptual framework in which the bacterial envelope functions as a spatially organized adaptive system rather than a passive permeability barrier. Under hypoxia, the mycobacterial cell envelope undergoes coordinated yet opposing remodeling trajectories, with inner membrane rigidification occurring alongside outer membrane fluidization. This organization generates a compartmentalized architecture that appears optimized for distinct functional requirements, including preservation of membrane-associated bioenergetic processes, maintenance of envelope plasticity, and attenuation of antibiotic interactions. By integrating membrane biophysics with layer-resolved lipidomics, we reveal that adaptive membrane responses occur with remarkable spatial precision across envelope layers. These findings extend current models of bacterial stress adaptation beyond bulk lipid remodeling and establish asymmetric envelope remodeling as an emergent principle of membrane organization. Given the prevalence of multilayered cell envelopes among actinomycetes and other medically important bacteria, this mechanism may represent a broader strategy by which microbial pathogens balance environmental adaptation, persistence, and antimicrobial resilience

## Conclusion

In summary, this study demonstrates that adaptation to hypoxia in *Mycobacterium smegmatis* is associated with extensive modulation of the mycobacterial cell envelope at both the biophysical and lipidomic levels. Hypoxic cells exhibited reduced growth, lower metabolic activity, and a significant decrease in whole-cell elastic modulus, indicating that oxygen limitation alters envelope mechanical properties. By integrating AFM nanoindentation with membrane biophysical analyses of isolated lipid fractions, we found that the two membrane layers respond differently to hypoxic stress: the inner membrane displayed increased lipid order and reduced hydration, whereas the outer membrane exhibited increased disorder and fluidity.

Layer-resolved lipidomics revealed that these biophysical changes were accompanied by membrane-specific shifts in lipid class abundance, acyl chain saturation, chain length distribution, and lipid molecular geometry. The enrichment of more saturated lipid species in the inner membrane and increased unsaturation within the outer membrane is consistent with, and may contribute to, the contrasting membrane states detected by Laurdan spectroscopy, fluorescence lifetime measurements, and FLIM. Notably, the emergence of distinct ordered and disordered domains within the hypoxic inner membrane suggests that hypoxia-induced membrane remodelling is not uniform but involves spatial reorganization of lipid populations.

Our findings further indicate that these envelope alterations are associated with functional changes in permeability and antibiotic response. Hypoxic bacteria showed reduced DAPI uptake and attenuated susceptibility to rifampicin, while membrane partitioning experiments suggested that altered interactions of rifampicin with hypoxic inner membrane lipids may contribute to this phenotype. Together, these observations support a model in which differential and remodelling of the inner and outer membrane layers enhances envelope asymmetry and alters membrane organization across multiple spatial scales, potentially influencing the permeability properties of the hypoxic cell envelope.

In conclusion, our findings reveal that hypoxia induces a spatially coordinated remodelling program across the mycobacterial envelope that links lipid composition, membrane organization, and antibiotic accessibility. Distinct adaptive responses within the inner and outer membrane layers generate a functionally asymmetric architecture characterized by inner membrane ordering and outer membrane fluidization. This organization is associated with reduced permeability and diminished rifampicin-membrane interactions, providing a mechanistic framework connecting oxygen limitation to antibiotic tolerance. More broadly, these results identify asymmetric envelope remodelling as a fundamental principle of bacterial membrane adaptation, demonstrating that individual membrane layers can be independently rewired to collectively optimize cellular fitness under stress. We propose that dynamic regulation of membrane asymmetry represents an underappreciated dimension of bacterial physiology with potential implications for persistence, pathogenesis, and antimicrobial intervention.

## Materials and Methods

### Materials

*M. smegmatis mc^2^155* (ATCC # 700084) was a humble gift from Dr. S. Chopra’s lab (CDRI, India). Middlebrook 7H9 broth was from BD Difco; BSA fraction V from MP Biochemicals, USA; Dextrose, catalase, and NaCl were purchased from SigmaAldrich. Chloroform, methanol and heptane of spectroscopic grade were purchased from Spectrochem. Tyloxapol and anhydrous glycerol were bought from Merck and EMPARTA, respectively. All of the products were used without further purification. Water used for aqueous buffer solutions was from a Millipore water purification system. Sulfosuccinic acid 1,4-bis (2-ethylhexyl) ester sodium salt (AOT), Chloroform, Methanol, Silica, Anthrone, TLC plates, Waters column

### Culturing *Mycobacterium smegmatis* under Normoxic and Hypoxic Conditions Normoxic condition

*Msm* (mc^2^155) bacteria were cultured in Middlebrook 7H9 media supplemented with 10% (v/v) albumin-dextrose-catalase, 10ml of 50% glycerol, and 2.5ml of 20% Tyloxapol under shaking conditions (120 rpm) at 37°C with aeration. Cells were harvested at 0.4 OD_600_ after 5-6 hours from 0.04 OD_600_.

### Hypoxic condition

*Msm* (mc^2^155) bacteria were cultured in Middlebrook 7H9 media supplemented with 10% (v/v) albumin-dextrose-catalase, 10ml of 50% glycerol, and 2.5ml of 20% Tyloxapol under shaking conditions (120 rpm) at 37 °C. Cells were harvested at 0.4 OD_600_. *M. smegmatis* was revived from glycerol stock kept in -80 °C in media and grown at 37°C, 120 rpm for 12 hours (primary culture). Bacteria were sub-cultured (initial OD 600 ∼ 0.05) from primary culture and grown until the mid-log phase (OD 600 ∼ 0.6-0.8) (approx. time- 8-9 hrs). The mid-log-phase bacterial culture was then adjusted to an OD600 of 0.04 (in media). Vacutainer tubes (Becton Dickinson, cat. no. 367812) with 4mL culture were used to optimize the hypoxic condition.

The tubes were kept in shaking at 37°C to allow for self-generation of hypoxia in the cultures. Fading or decolorization of methylene blue (added at final concentration of 1.5 mg/L) was used to detect once hypoxia is attained. For scaling up to higher volumes of bacterial culture for lipid extraction, roller bottles (1L capacity) were used and media was added in a way so that 0.5 HSR (headspace ratio) was maintained. In a 1L capacity roller, 800ml of bacteria can be cultured maintaining the 0.5 HSR. Hypoxia cultures were kept in shaking at 120rpm, completely sealed at 37°C for 30 hours. Cells were harvested at 0.4 OD_600_.

### Lipid extraction

Lipids were extracted selectively from the outer and inner membrane layers of *Msm* using a previously reported method. For the recovery of all non-covalently bound OML lipids, 1 mL of reverse micellar solution (RMS; 10 mM sulfosuccinic acid 1,4-bis (2-ethylhexyl) ester sodium salt (AOT) in heptane) was added to every 10 mg dry weight of cells. After the first overnight extraction, two re-extractions of 15–30 min each under continuous stirring conditions at room temperature (RT). For the extraction of IML lipids, the RMS-extracted cells were thoroughly washed with water, dried, and then extracted with chloroform: methanol: water (C:M:W::2:1:0.1). After an overnight extraction, two more extractions were done for 30 min each, where, for each 10 mg of dry cell mass, 3 ml of CMW was used. All the extractions were carried out under continuous stirring on a magnetic stirrer at RT. The lipids were recovered from the supernatants, filtered, and the solvent was removed, and further purified (Adhyapak et al., 2020).

### Recovery of the AOT-free fraction of lipids

To remove AOT contamination from the lipid mixture for further studies, purification of the OML from the mixture was done using column chromatography packed with silica mesh with a mobile phase of gradient methanol in chloroform. Almost all lipids were eluted, which was confirmed from the fractions run on thin-layer chromatography (TLC) plates developed with 1% anthrone spray.

### Characterisation of lipids

Shaded ellipses represent group clustering distributions for biological replicates. The percentage variance explained by each principal component is indicated on the corresponding axes. Together, these analyses demonstrate that hypoxia induces membrane layer–specific lipidomic reprogramming in both the inner and outer membrane systems

### LCMS

#### Sample preparation

Lipid samples namely IML Normoxic and Hypoxic were taken in duplicates for one run at a time at a final concentration of 4mg/mL. Reserpine was taken as the internal standard (IS) for quantification of the identified lipids. Standard lipids were also taken as controls to check its presence by the instrument. ^25,34^

#### LC

Extracted lipids were then analysed in both positive and negative ion mode using a 1290 Infinity UHPLC System attached to a 6550 iFunnel Q-TOF system (Agilent Technologies, USA). The lipids were identified using databases available online. An Agilent 1200 HPLC (Agilent Technologies; Palo Alto, CA) with a 2.1 inner diameter (ID) × 150 mm, 3.5 μm XBridge C18 column (Waters Corp.; Milford, MA) heated to 45°C was used with a binary solvent system and a flow rate of 0.175ml/min. ^25,34^

#### MS

An Agilent Mass Spectrometer Quadrupole time-of-flight (MS Q-TOF) G6545XT equipped with a Dual AJS ESI mode source was used for accurate mass analysis of the LC eluent. Positive- (+) and negative- (-) ion data were generated by operation of the mass spectrometer with a capillary voltage of 4000 V, nebulizer of 45 psig, drying gas of 8 l/min, gas temperature of 300°C, fragmentor of 175 V, charging voltage of 2000 V, skimmer of 65 V, and octopole radio frequency voltage of 750 V. Mass spectra were acquired at a rate of 1.00 spectra/s and data were collected as profiled spectra over a mass range of 100 to 3,200 Da. Mass calibration was performed with an Agilent tune mix from 100 to 3,200 Da.

Data were collected with the Agilent Qualitative Analysis of MassHunter Acquisition Data software, version 10.0. Positive-ion mass spectra were acquired in Auto MS/MS mode, and collision energies with a slope of 6.5 V/100 Da and an offset of 2.0 V were used for fragmentation.

### Quantification of lipids

These lipids were quantified using the information processed from the MassHunter software ^25^. Area and molecular weight of the lipids were used for quantifying the mol% of a lipid present in a membrane. Reserpine was used as the internal standard for quantification purposes. ppm and molarity were calculated to ultimately calculate the mol% of each lipid present.

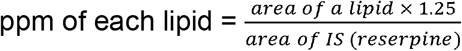

where, 1.25 is the ppm (concentration) of reserpine (internal standard) used.

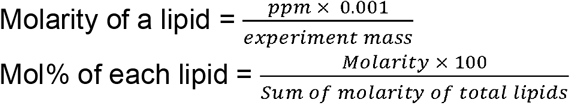

### Liposome preparation

Extracted lipids were dissolved in chloroform and stored at −20°C until further use. Lipid films were made by adding the desired lipid in chloroform to get a final concentration of 0.25 mg/mL and the probe at 0.00125 mg/mL concentration to form a final lipid concentration at a probe/lipid ratio of 1:200. The organic solvent was dried with a gentle stream of N2 gas followed by overnight lyophilization. Lipid suspensions were prepared by gentle hydration and the freeze−thaw method. The lipid films were hydrated using Tris-MgCl2 buffer at a temperature of 55-60°C, where after initial sonication for 10 min, the samples were subjected to 5 freeze-thaw cycles for experiments to yield large unilamellar vesicles (LUVs).

### Laurdan Generalised polarisation (G.P.) spectroscopy

The steady-state fluorescence spectroscopic measurements were performed on a Horiba FluoroMax Plus (RM360), which has a temperature controller. Laurdan, a solvatochromic probe, was employed to determine the membrane order. A final lipid concentration of 0.25 mg/mL with a probe/lipid ratio of 1:200 was used. Laurdan was excited at 350 nm, and the emission was measured from 370 to 580 nm, considering a temperature range from 5°C - 80°C (accuracy of ± 0.1°C) with 3 minutes of equilibration

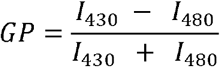

where I430 and I480 refer to the intensities at 430Onm and 480Onm, which are characteristic for an ordered (gel) lipid phase and fluid liquid-crystalline phase, respectively. The GP values range from +1 to −1.

### Giant unilamellar vesicles (GUVs)

Lipids were mixed to obtain a final concentration of 4 mg/mL with Laurdan at the ratio of 1:200. Giant unilamellar vesicles (GUVs) were prepared using the electroformation method in a temperature-controlled custom-made chamber using optically transparent and electrically conductive indium tin oxide-coated glass coverslips (SPI Supplies, West Chester, PA). Lipid mixture (100 µL) was spin-coated (1000 rpm for 20 s) onto the coverslips and subsequently dried under vacuum. The coverslips were placed within the electroformation cell, and the cell was filled with Tris-MgCl_2_ buffer. Lipids were hydrated within the cell at a temperature of 65°C, and a low-frequency alternating current field (sinusoidal wave function with a frequency of 10 Hz and an amplitude of 2Vp-p) was applied for 120 min. The cell was gradually cooled to RT before imaging the GUVs. The GUVs were then imaged under FLIM, and the images collected were of 512 × 512-pixel resolution.

### Fluorescence lifetime imaging microscopy (FLIM)

FLIM was performed on a MicroTime 200 (Picoquant, Gmbh, Germany) using Time-tagged Time-resolved (TTTR) methodology. This setup is attached to an inverted microscope (IX71) equipped with a water immersion objective (UPlan SApo NA 1.2, 60X, WD = 0.28 mm). Laurdan in GUVs were excited at 405 nm using a pulsed diode laser. Laurdan fluorescence emission was collected using a 460/60 bandpass filter. The collected fluorescence signal was directed to a single photon avalanche photodiode (SPAD) detector. The images were acquired with 512 x 512-pixel resolution. The fluorescence decays were analyzed using the inbuilt software of MicroTime 200 and were fitted using iterative deconvolution, to bi-exponential function.

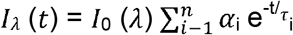

where I(t) is the intensity of the fluorescence at time t, *α*_*i*_ is the pre-exponential factor for the fraction of the fluorescence intensity, τ_*i*_ is the fluorescence lifetime of the emitting species, and n is the number of exponentials used. The average fluorescence lifetime was calculated using the following relation.

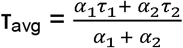

where α1 and α2 are the pre-exponential factors representing the fractional contribution of the decaying component with a lifetime τ_i_.

Fluorescence lifetime refers to the average duration that a fluorophore remains in its excited state before returning to the ground state through either radiative processes (emitting a photon) or non-radiative mechanisms, such as thermal vibrations. Employing time-resolved fluorescence lifetime imaging microscopy (FLIM), I collected fluorescence decay data from the fluorescent probe at two distinct emission wavelengths. I fitted the experimental data using nonlinear least squares methods to obtain the fluorescence lifetime, which is calculated using the following formula:

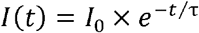

The parameter *I(t)* represents the fluorescence intensity at time t, *I_0_* denotes the initial fluorescence intensity, and τ signifies the fluorescence lifetime.

### Time-Correlated Single-Photon counting (TCSPC)

Fluorescence decay was recorded using a DeltaFlex time-correlated single-photon counting (TCSPC) system (HORIBA Scientific). Samples were excited at 360 nm with Titanium Sapphire laser source. Emission decays were measured at 430 nm at magic angle (54.7°) polarization. All measurements were performed at 23±1°C. Fluorescence decay traces were fitted to triexponential or sum of exponential functions using iterative deconvolution in EzTime software.

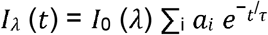

where τ_i_ and a_i_ are the lifetime and amplitude of the *i^th^*component, respectively.

### Atomic Force Microscopy

AFM Imaging was carried out in Intermittent contact (AC) mode and force spectroscopy was done in contact mode with MFP-3D atomic force microscope (Asylum Research, Santa Barbara, CA, USA.) Bacteria were grown under normoxic and hypoxic conditions. The bacteria were centrifuged and pelleted down. It was resuspended in PBS to give a wash that would wash off excess media component. Centrifuge it again and then resuspend in PBS. Add approximately 10-15 µL of bacterial sample onto a clean glass slide. Give a gentle wash using water and let it dry. Then image under AFM dry mode.

### Flow Cytometric Assessment of Membrane Integrity and Bacterial viability Following Hypoxia and Drug Exposure

Bacterial cultures were subjected to normoxic or hypoxic conditions as described above to establish replicating and oxygen-limited states, respectively.

DAPI was used as a probe to assess membrane integrity. After culturing the bacteria under the respective condition, i.e., normoxic and hypoxic, DAPI was added to it at a concentration of 1µg/mL for 30 mins at 37°C in the dark. The bacteria were then washed and resuspended in PBS. Data acquisition was done using a flow cytometer equipped with a 405 nm excitation laser. A total of 50,000 events were recorded per sample. Forward and side scatter parameters were used to gate the bacterial population and exclude debris. Known permeabilizers, Triton-X 100 (TX100) and lysozyme, were used for positive controls. After normoxic and hypoxic bacterial culturing, bacteria was treated with TX100 at a concentration of 0.1% for 5 mins and then lysozyme at a concentration of 2mg/mL for 30 mins at 37°C. The bacteria were then washed. DAPI was then added to it at a concentration of 1µg/mL for 30 mins at 37°C in the dark. The bacteria was then washed, resuspended in PBS and then taken for acquisition. Untreated bacteria served as a negative control to assess baseline DAPI uptake in normoxic and hypoxic *Msm.* Positive control consisting of known membrane permeabilizers Triton-X 100 and Lysozyme, was used to determine the extent to which deliberate envelope permeabilization could enhance DAPI uptake under each condition.

Following adaptation, cultures were exposed to rifampicin or metronidazole for 24 h at concentrations based on their respective minimum inhibitory concentrations (MICs). Rifampicin was used at 2, 8, 32, and 128 µg/mL, while metronidazole was used at 20, 100, 400, 1600, and 6400 µg/mL. After drug treatment, bacterial cells were pelleted down. Pellets were washed and resuspended in sterile phosphate-buffered saline (PBS), and cell density was adjusted to an OD_₆₀₀_ of 0.2 to ensure uniform distribution across samples.

To assess bacterial viability, cells were stained with propidium iodide (PI) at a final concentration of 3 µg/mL and incubated for 5 min at 37°C in the dark. Following staining, cells were pelleted again under the same centrifugation conditions, washed once with PBS to remove excess dye, and finally resuspended in PBS for flow cytometric acquisition.

Data acquisition was performed using a flow cytometer equipped with a 561 nm excitation laser. A total of 50,000 events were recorded per sample. Forward and side scatter parameters were used to gate the bacterial population and exclude debris. Heat-killed bacteria were used as a positive control to define the PI-positive population and establish the membrane-compromised gate. The percentage of PI-positive cells was quantified and used as a measure of membrane damage and loss of viability following drug exposure under normoxic and hypoxic conditions. All analyses were performed using standard flow cytometry software, and PI-positive percentages were used for comparative statistical analysis.

### Determination of Partition Coefficient (Kp)

The partition coefficient (K_p_) of rifampicin between the lipid membranes and the aqueous buffer was determined using a UV-visible spectrometry technique. Briefly, 10 μM rifampicin was added to increasing concentration (0-1000 μM) of lipid vesicles and incubated at 37□ C for one hour and the absorption spectra were recorded using a multi-detection microplate reader (ThermoFisher) from 200-600 nm with 1 nm intervals at 37 □ C. The intensities were then processed to obtain the third derivative with respect to the wavelengths using the nprot Kp calculator. For the calculation of Kp values, third derivative intensities were considered by fitting the experimental data to the following equation by the nonlinear regression method using Origin 2021a.

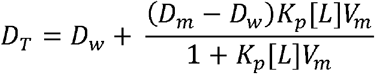

where D represents the third derivative intensity

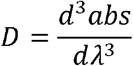

obtained from the absorbance of the total rifampicinconcentration (D_T_), rifampicin distributed in lipid membrane phase (D_m_) and rifampicin distributed in the aqueous phase (D_w_); [L] is the molar lipid concentration and V_m_ is the molar volume derived earlier for each lipid mixtures.

## Acknowledgments and Funding

This work was supported by the DBT/Welcome Trust India Alliance Intermediate Fellowship (IA/I/21/1/505624) awarded to SK. Central facilities at CSIF, IIT Bombay are gratefully acknowledged. We would also like to acknowledge Dr Aswin Srivatsav for the MATLAB program used for lipidomics.

