## Supplementary Information for "Hypoxia drives asymmetric envelope remodelling to promote membrane adaptation and antibiotic tolerance in mycobacteria"

**Establishment of the Hypoxic Culture Model and Assessment of Bacterial Viability**

Hypoxia was induced in a sealed standing culture model. Methylene blue at a concentration of 1.5 µg mL^-1^ was added to the culture medium as a redox indicator to monitor oxygen availability.

**Normoxic**

**O_2_ present**

**Intermediate**

**After 20 hrs**

**Hypoxic After ~30 hrs.**

**Low O_2_**

***Supplementary Figure S1.*** *Representative photographs showing the progressive establishment of hypoxia in Mycobacterium smegmatis cultures. Methylene blue served as a visual indicator of oxygen availability, remaining blue under normoxic conditions and becoming colorless upon oxygen depletion. Images represent normoxic (left), intermediate (~20 h; middle), and hypoxic (~30 h; right) cultures.*

The dye stays blue under normoxic conditions because of the dissolved oxygen. During incubation, the dye decolourised as oxygen was progressively consumed, showing the establishment of hypoxic conditions. Supplementary Figure S1 shows representative images of the cultures under normoxic, intermediate (20 h), and hypoxic (~30 h) conditions.

***Supplementary Figure S2.*** *Mycobacterium smegmatis grown under normoxic (left) and hypoxic (right) conditions. Hypoxic cultures exhibited a markedly reduced number of colonies compared to normoxic cultures, consistent with decreased bacterial replication under oxygen-limited conditions.*

Bacterial viability was determined by colony-forming unit (CFU) counting to confirm the physiological state of the cultures after induction of hypoxia. Aliquots of normoxic and hypoxic cultures were serially diluted in phosphate-buffered saline (PBS), plated on Middlebrook 7H10 agar supplemented with OADC and incubated at 37°C until visible colonies formed. Representative plates are in Supplementary Figure S2. The low number of colonies recovered from hypoxic cultures compared to normoxic cultures confirmed that the bacteria were in a low-metabolic, non-replicating state and remained viable.

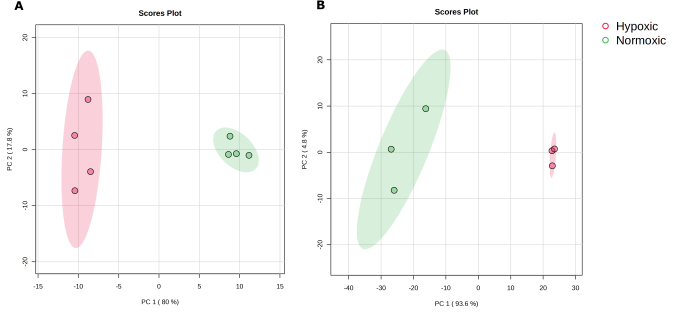

**Supplementary Figure S3.** Principal component analysis reveals distinct hypoxia-induced lipidomic remodelling in inner and outer membrane systems. Principal component analysis (PCA) score plots generated from lipidomic datasets of inner membrane lipids (IML) (A) and outer membrane lipids (OML) (B) isolated from normoxic and hypoxic M. smegmatis cultures, reflecting extensive compositional reorganization of the outer and inner membranes under oxygen limitation.

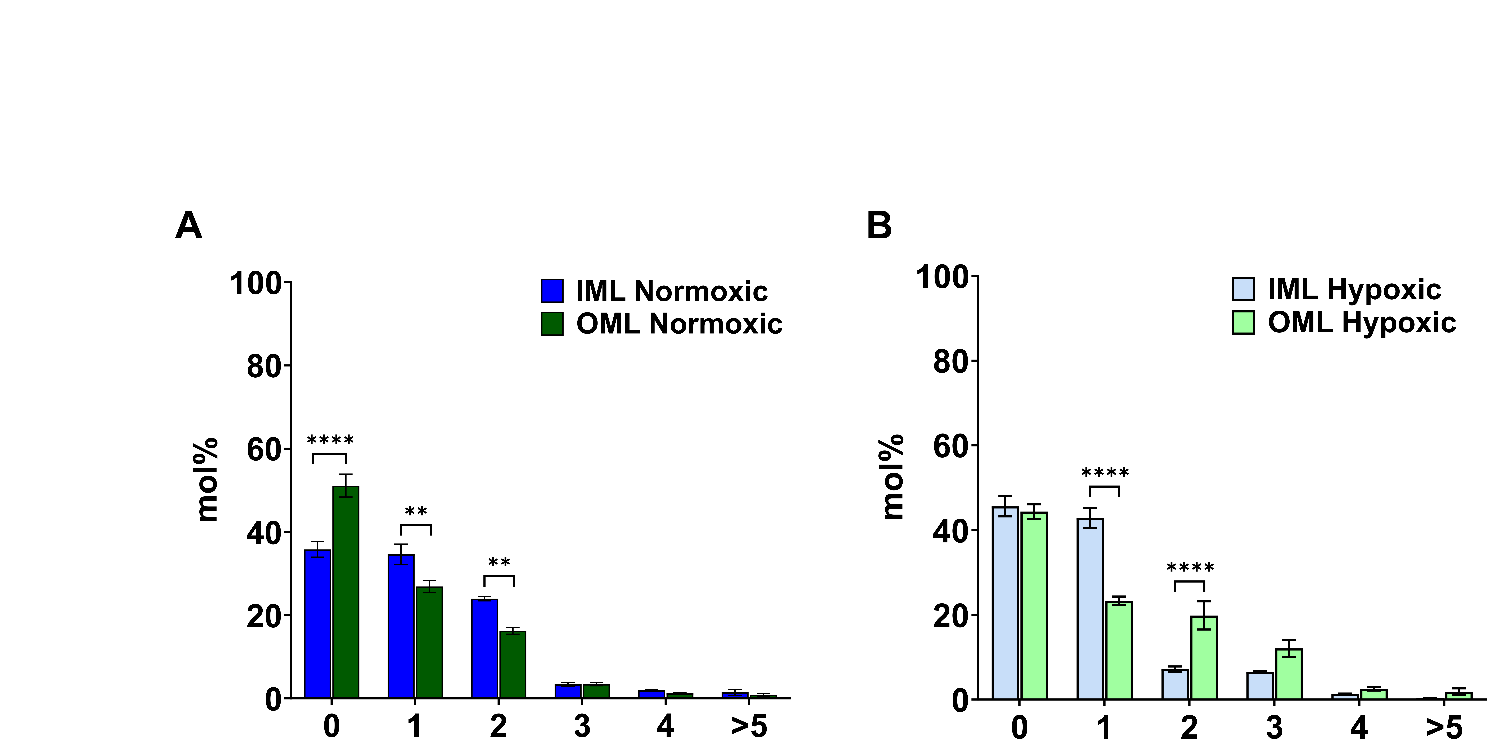

***Supplementary Figure S4.*** *Lipidomic profiling of membrane lipid unsaturation in the inner and outer membrane fractions of Mycobacterium smegmatis under normoxic and hypoxic conditions. Comparison of lipid unsaturation between IML and OML under (A) normoxic and (B) hypoxic conditions.*
